# Subliminal risk influences subjective value in the ventromedial prefrontal cortex

**DOI:** 10.1101/2025.03.11.642695

**Authors:** Patrícia Fernandes, Sangil Lee, Joseph W. Kable, Philipp Seidel, Jorge Almeida, Johan Eriksson, Bruno de Sousa, Fredrik Bergström

**Affiliations:** CINEICC, Faculty of Psychology and Educational Sciences, University of Coimbra, Portugal; Helen Wills Neuroscience Institute, University of California, Berkeley, CA, USA; Department of Psychology, University of Pennsylvania, Philadelphia, PA 19104, USA; Department of Psychiatry and Psychotherapy, Tübingen Center for Mental Health, University of Tübingen, Germany; Umeå center for Functional Brain Imaging (UFBI); Department of Psychology, Umeå University, Sweden; Department of Psychology, Gothenburg University, Sweden

**Keywords:** subliminal, risk, subjective value, conscious awareness, fMRI

## Abstract

The relationship between conscious awareness and decisions has been heavily debated. Here we investigated whether subliminal probabilities are integrated with conscious rewards to form subjective value (SV) representations in the anterior ventral striatum (aVS) and ventromedial prefrontal cortex (vmPFC). Participants played an incentivized competitive game with risky choice to accumulate points across trials in a behavioral and fMRI experiment. The game was a modified attentional-blink paradigm that rendered a probability cue unseen (indicating a 100% or 0% chance to win a risky reward). Following the probability cue, participants chose between a safe (1 point with certainty) or risky option (>1 or 0 points depending on probability cue). The risky reward was either 2 or 5 points, varying across trials. In some trials the probability cue was absent (replaced by a random distractor) and the probability to win the risky reward was 50%. When probability cues were unseen, they did not influence choice, as value-maximizing choice (d’) was not greater than chance, but they did influence reaction time in both experiments. Consistent with SV integration, the BOLD signal in aVS and vmPFC was higher for both conscious rewards (high > low) and subliminal probabilities (high > low) and could not be explained by subliminal salience (cue present > absent). Moreover, multivariate pattern similarity between conscious rewards and subliminal probabilities in vmPFC suggest integration into an abstract value representation. Additionally, we found brain-wide subliminal probability and salience effects. Taken together, these results suggest that conscious awareness is not necessary for probability to be integrated with conscious rewards to form an abstract “common currency” SV representation in vmPFC. Additionally, brain-wide subliminal probability and salience effects suggests information can have “global access” without conscious awareness.

---

Our intuition is that we are consciously aware of the information influencing our value-based decisions. However, this intuition may be misleading as we are only consciously aware of a fraction of all neural processes at any given time. Whether information outside conscious awareness can influence our decisions has been heavily debated (for critical review, see Newell & Shanks, 2014). However, the few existing neuroimaging studies about subliminal influences on value-based decisions have not shown subliminal value processing outside of subcortical areas. Here we used subliminal probability cues to investigate whether they were integrated with conscious rewards to form subjective value (SV) representations not only in the anterior ventral striatum (aVS) but also in the ventromedial prefrontal cortex (vmPFC).

When consciously aware of choice information, we typically select the option with the highest SV. In the context of risk (i.e., uncertain outcomes with known probabilities), SV is derived by discounting the reward value according to the probability of not receiving it. Choice-related information, such as probabilities and rewards, is thought to be integrated into abstract domain-general (“common currency”) SV representations in the aVS and vmPFC (Bartra et al., 2013; Chib et al., 2009; Dang et al., 2024; Kable & Glimcher, 2009; Kobayashi & Hsu, 2019; D. J. Levy & Glimcher, 2012; I. Levy et al., 2010; McNamee et al., 2013; Pegors et al., 2015; Shuster & Levy, 2018). This neural SV signal in aVS and vmPFC has been dissociated from salience, which follows a U-shaped relationship with SV – i.e., higher BOLD signal for both high rewards and high punishments (Bartra et al., 2013). Instead, salience has been linked to brain areas such as the anterior insula (aINS), caudate nucleus, dorsomedial prefrontal cortex (dmPFC), dorsolateral prefrontal cortex (dlPFC), and inferior posterior parietal cortex (PPC) (Bartra et al., 2013; Kahnt & Tobler, 2017; Litt et al., 2011; Zink et al., 2004). However, the relationship between conscious awareness, SV, and salience remains largely unexplored.

Studies that explored subliminal influences on value-based decisions have shown that subliminal rewards can be processed subcortically and influence behavior. Specifically, subliminal reward cues have been shown to influence ventral pallidum and grip force (Pessiglione et al., 2007), bilateral aVS and instrumental learning (Pessiglione et al., 2008), and cognitive task performance (Aarts et al., 2008; Bijleveld et al., 2009, 2012a; Custers & Aarts, 2010). Based on these empirical findings, it was suggested that subliminal value processing is limited to rudimentary subcortical processing, while conscious awareness is necessary for “full” value processing in the prefrontal cortex (Bijleveld et al., 2012b).

The importance of the prefrontal cortex for conscious awareness underpins prominent theories of consciousness, such as the Global Neuronal Workspace (GNW) (Dehaene & Changeux, 2011; Dehaene & Naccache, 2001; Mashour et al., 2020) and Higher-Order Thought (HOT) theory (Lau & Rosenthal, 2011). However, these theories have conceded that subliminal information can influence the prefrontal cortex to a lesser extent than conscious information (Lau & Rosenthal, 2011; Melloni et al., 2023). Indeed, it has been shown that subliminal information can engage dlPFC and associated functions (Bergström & Eriksson, 2014, 2018; Lau & Passingham, 2007; van Gaal et al., 2010). Moreover, it seems that SV is automatically computed in vmPFC for consciously perceived stimuli in the absence of explicit valuation tasks (Lebreton et al., 2009; Levy et al., 2011; Tusche et al., 2010). Given these considerations, we hypothesize that SV computations can occur without conscious awareness in aVS and possibly also in vmPFC.

Here we used a novel attentional blink paradigm with trial-wise measures of awareness and embedded risky choice task to investigate the relationship between conscious awareness and neural representations of SV and salience. In two experiments (a behavioral and an fMRI), participants were presented with subliminal probability-cues prior to choosing between safe (1 point with certainty) or risky options (>1 or 0 points). Subliminal probability cues were either associated with 100% or 0% chance of winning the risky reward and 50% when the cue was absent. The risky reward was either 2 or 5 points and consciously presented at the time of choice. Our goal was to see whether subliminal probabilities were integrated with conscious rewards in a way consistent with SV. Behaviorally, we expected reaction time to be faster for higher rewards and higher probabilities. Neurally, we expected BOLD signal in aVS and vmPFC (i) to be higher for both conscious rewards (high > low) and subliminal probabilities (high > low), suggesting SV integration; (ii) not to be explained by subliminal salience (probability cue presence > absence); (iii) to exhibit pattern similarity between conscious rewards and subliminal probabilities, suggesting a shared neural code; and (iv) to correlate negatively with reaction time. We found that BOLD signal in the vmPFC exhibited all characteristics consistent with SV, while subliminal salience was found in brain areas previously associated with (conscious) salience.

## Materials and Methods

### Participants

For the behavioral experiment, we recruited 42 healthy volunteers from the University of Coimbra campus area. All participants had normal or corrected-to-normal vision, gave written informed consent, and received extra course credits for participation. The participant with highest score in the choice game additionally received a 50-euro reward voucher. Participants were excluded for not following instructions (n = 3), not having any trials in a condition necessary for the analyses (i.e., never choosing the risky option when reward was low; n = 4), or having unusually high value-maximizing choice (d’) score when probability cue was reported unseen (n = 1; d’ > 3 SD; to avoid the possibility of conscious influences). We therefore used 34 participants (27 females; 18 – 44 age range, M = 21, SD = 6 years) for our analyses.

For the fMRI experiment, we recruited 57 healthy volunteers from the University of Coimbra campus area and through social media. All participants had normal or corrected-to-normal vision, gave written informed consent, received extra course credits (if applicable), and a 20-euro reward voucher for participation. The participant with highest score in the choice game additionally received a 50-euro reward voucher. Participants were excluded prior to the fMRI experiment for seeing the probability cue on more than 20% of practice trials (n = 15) and excluded after the fMRI experiment for not having any trials in a condition necessary for the analyses (never choosing the risky option when reward was low; n = 4), or having unusually high value-maximizing choice (d’) score when probability cue was reported unseen (n = 2; d’ > 3 SD; to avoid the possibility of conscious influences), or loss of MRI data because of technical problems at MRI scanner (n = 3). We therefore used 29 participants (22 females; 18 – 41 age range, M = 22, SD = 6 years) for our analyses, except for the aVS region of interest analyses where a participant was excluded for not having signal in aVS. The study was approved by the Ethics Committee of the Faculty of Psychology and Educational Sciences of the University of Coimbra, Portugal.

### Stimuli and Procedure

The experiments were designed and executed with A Simple Framework (ASF; Schwarzbach, 2011) in MATLAB R2019a (MathWorks Inc., Natick, MA, USA) and presented at a 60Hz refresh rate on a Samsung SyncMaster for behavioral experiment and a BOLDscreen 32UHD (Cambridge Research Systems Ltd., Rochester, UK) for the fMRI experiment. We used a Cedrus RB-730 button box for the behavioral experiment and Cedrus Lumina button box for the fMRI experiment for accurate reaction time measures.

Both experiments were incentivized competitive risky choice games where participants were instructed to accumulate as many points as possible across all trials for the possibility to win a 50-euro reward voucher. The behavioral experiment had 480 trials across eight runs (60 trials, 40 with and 20 without probability cues, per run) and the fMRI experiment had 336 trials across four runs (84 trials, 56 with and 28 without probability cues, per run). The game was a modified attentional-blink paradigm embedded with a risky choice task (Figure 1). After an initial inter-trial-interval (ITI) with a fixation cross, there was a rapid serial visual presentation of several irrelevant abstract symbols (distractors) and two targets to identify (T1 and T2). T1 was five yellow “arrows” randomly pointing left or right superimposed on a random distractor, and the task was to discriminate the direction of most arrows (left or right). The T1 task was difficult on purpose to induce an attentional blink or deficit at the time T2 was presented. T2 was one of two abstract symbols associated with a 100% or 0% probability of winning a risky reward. Unbeknownst to the participants, there were also trials where T2 was absent and replaced by a random distractor, on these trials there was a 50% probability of winning the risky reward to maintain the illusion of constant T2 presence. At the end of each trial there were three queries. First, participants had to choose between a safe (1 point with certainty) or risky option (> 1 or 0 points depending on which probability cue was presented). The risky reward amount was either 2 or 5 points and varied across trials. The counterbalanced choice and reward information was consciously presented at the time of choice. Second, participants had to report their perceptual awareness of T2 (the probability cue) using a perceptual awareness scale (PAS; Sandberg et al., 2010). Third, they had to report the direction of most arrows (left or right). Each trial ended with feedback indicating if T1 was correct or not and how many points were won on that trial. The modified attentional-blink paradigm was designed to maximize the number of trials where T2 was reported unseen, while maintaining a relatively long (83 ms) presentation duration to maximize subliminal effects.

**Figure 1.**
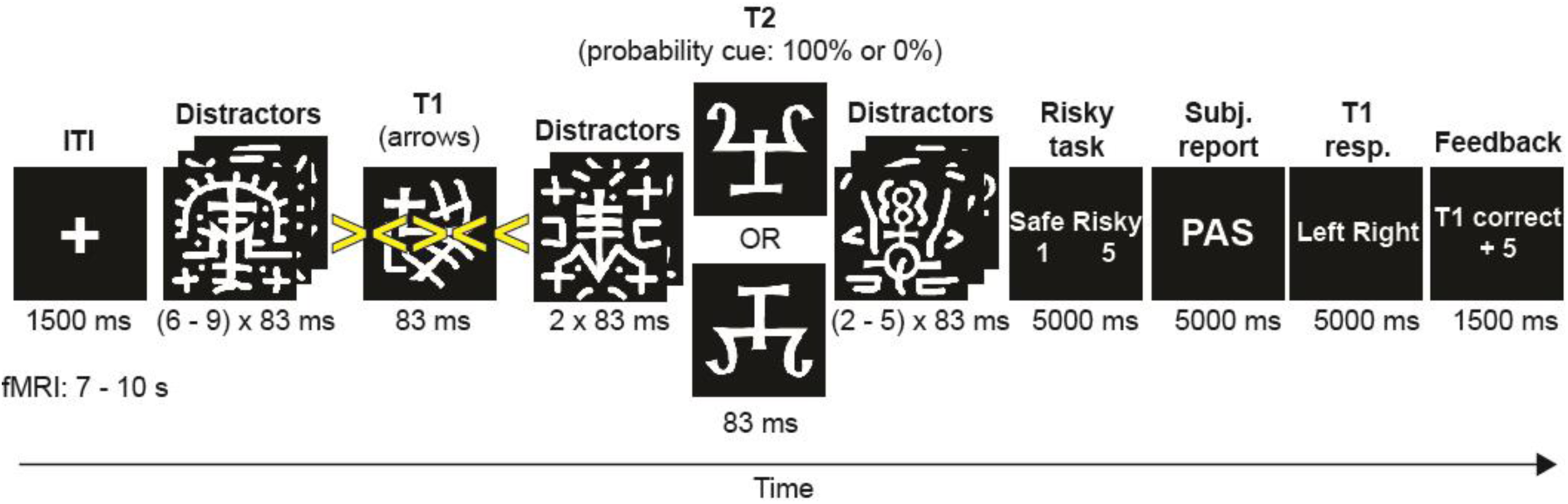
Trial procedure. Five yellow arrows (T1) and one of two abstract symbols (T2; probability cue) were presented in a rapid serial visual presentation. At the end of each trial, participants had to make a safe (1 point with certainty) or risky (>1 or 0 points depending on T2) choice, report their perceptual awareness of T2, and report where most arrows pointed (left or right). Response screens lasted until a button was pressed or until 5 s passed. The risky reward amount varied between 2 and 5 points across trials. On some trials T2 was absent and replaced by a random distractor. On T2 absent trials the risky reward was won on 50% of trials.

Participants were given a set of specific instructions to follow during the experiment. First, the goal was to accumulate as many points as possible across all trials to have the possibility to win a 50-euro reward voucher. Second, a correct response to the arrow direction (T1) was a prerequisite to get any points on a trial (an incorrect response automatically led to 0 points) to ensure that attention was primarily on identifying T1, which is necessary to create the attentional blink that renders T2 unseen (Martens & Wyble, 2010). Third, points did not depend on PAS ratings to not incentivize participants to report T2 as unseen when in fact it was seen. Fourth, prior to practice, participants were told that their total points would be adjusted based on their accuracy (i.e., d’) such that any systematic strategy (e.g., always guessing risky or safe) would reduce their total points and be counterproductive to winning the game. This was to incentivize participants to make a balanced number of safe and risky choices while relying on their “gut-feeling” to guess the most valuable guesses each trial, which is important because systematic strategies run the risk of making it more difficult to find subliminal behavioral effects.

The trial procedure was almost identical for the behavioral and fMRI experiments with two exceptions. First, the fixation duration was 1.5 s for the behavioral and 7 – 10 s for the fMRI experiment to account for the sluggishness of the BOLD signal. Second, the perceptual awareness scale used a 3-point scale (1 = no visual experience, 2 = vague visual experience, 3 = almost clear or clear visual experience) for the behavioral and a 2-point scale (1 = no visual experience, 2 = vague, almost clear, or clear visual experience) for the fMRI experiment because the MRI compatible button-box had two buttons. However, participants were trained on a 3-point PAS scale prior to the fMRI session to become accustomed to making perceptual distinctions between no, vague, or almost clear visual experiences of T2.

Participants read the instructions on a sheet of paper and a researcher further explained them in detail before they completed a practice session. The practice session initially used a slower presentation duration where T2 was consistently seen relatively clearly and later used the same presentation speed as the experiment to prepare the participants for the task and for the researcher to make sure participants understood and followed the instructions. Prior to the fMRI experiment, this practice session was used to filter out participants that saw T2 (PAS > 1) on more than 20% of trials. This was to ensure that participants in the fMRI experiment would have a high number of unseen trials for analyses. In the behavioral experiment, participants proceeded with the experiment session directly after the practice session. In the fMRI experiment there could be one week or more between practice and MRI session, so participants were reminded of the instructions again before entering the MRI scanner (but no active practice).

### MRI acquisition

MRI data were collected with a 3T MAGNETOM Vida whole-body MR scanner (Siemens Healthineers, Erlangen, Germany) at Hospital da Luz de Coimbra initially with a 20-channel head coil (n = 7) and later when available a 32-channel head coil (n = 22). MRI acquisition occurred in one session with an anatomical image acquired between four functional runs (2260 volumes in total). Anatomical MRI images were acquired using a T1-weighted magnetization-prepared rapid gradient echo (MPRAGE) sequence [repetition time (TR) = 2300 ms, echo time (TE) = 2.32 ms, voxel size = 0.9 x 0.9 x 0.9 mm^3^, flip angle = 8 degrees, field of view (FoV) = 240 x 240, matrix size = 256 x 256, bandwidth (BW) = 200 Hz/px, GRAPPA acceleration factor 2]. Functional MRI (fMRI) data were acquired using a T2*-weighted gradient echo planar imaging (EPI) sequence (TR = 2000 ms, TE = 36 ms, voxel size = 3 x 3 x 3 mm^3^, slice thickness = 3 mm, FoV = 210 x 210, matrix size = 70 x 70, flip angle = 75 degrees, BW = 1786 Hz/px). Each image volume consisted of 50 contiguous transverse slices recorded with interleaved slice order oriented parallel to the line connecting the anterior commissure to the posterior commissure and tilted 30 degrees for better signal acquisition in the vmPFC.

### Preprocessing of MRI data

MRI data were preprocessed with fMRIPrep 23.2.0a2. See below for a summary of the preprocessing pipeline and the supplemental material for a more detailed fMRIPrep boilerplate. The T1-weighted anatomical image was corrected for intensity non-uniformity, used as T1-reference in the workflow, skull-stripped, tissue segmented (grey matter, white matter, and CFS), and spatially normalized into MNI space using non-linear registration. For functional EPI images the following steps were done. A reference volume was generated with custom fMRIPrep methodology for use in head-motion correction. The EPI images were head-motion corrected using six rotation and translation parameters and slice-time corrected. The BOLD reference was co-registered to the T1-reference. Several confounding time-series were estimated based on the preprocessed BOLD series and 17 were later used as covariates in the General Linear Models (GLMs). The preprocessed BOLD was normalized to MNI space, and 8 mm Gaussian FWHM filter was applied with SPM12 prior to analysis.

### Statistical analysis

#### Behavioral analysis

Since the purpose of the T1 task was to be difficult enough to induce a strong attentional blink effect on T2, we included trials irrespective of whether T1 was answered correctly or not in our analyses to maximize statistical power. To assess subliminal behavioral effects, we only used trials where T2 was reported as unseen (PAS = 1) and removed trials if risky choice task reaction time was < 200 ms as these trials would likely be caused by accidentally pressing the button before processing choice information. “Hits” were defined as risky choice when probability was 100% to win risky reward, “false alarms” (FAs) as risky choice when probability was 0%. Value-maximizing choice (d’) was defined as z(hits) minus z(FAs). The criterion was calculated by multiplying the sum of z(hits) and z(FAs) by −0.5. The d’ score was used to measure direct influences, criterion was used to measure choice bias, and reaction time to measure indirect influences on behavior. Reaction times were log10-transformed prior to statistical analyses.

#### fMRI analysis

##### General Linear Model (GLM)

Similar to the behavioral analysis, we included trials irrespective of whether T1 was answered correctly or not in our analyses to maximize statistical power. For within-participant modeling with SPM12, we used a GLM with restricted maximum likelihood estimation and six regressors of interest (Rew5Prob1, Rew2Prob1, Rew5Prob0, Rew2Prob0, Rew5abs, Rew2abs), an “other” regressor with trials where probability cues were reported seen, and 44 motion/noise parameters (24 head motion parameters, 5 cerebral spinal fluid CompCor components, 14 white matter CompCor components, and a frame-wise displacement parameter). We equalized the number of CompCor components across participants and runs by selecting the highest number of components that all subjects had available. Regressors of interest were modeled from the onset of the probability cue until 300 ms after the onset of the risky choice task screen regardless of reaction time and were convolved with the canonical hemodynamic response function in SPM12. The high-pass filter had a cut-off at 128 s and the autocorrelation model was global AR (1).

##### Contrasts

The GLM was used compute beta maps for three group-level contrasts: (i) high > low risky rewards (5 > 2 points), (ii) high > low probability to win risky reward (100% > 0%), and (iii) high > low salience (probability cue present > absent). The salience contrast is thus not influenced by reward or probability because the average reward and probability values are the same when cue is present and absent.

##### Univariate analysis

We used the group-level whole-brain contrasts to look for significant BOLD signal change in voxels within a group-level grey matter mask, and used Threshold-Free Cluster-Enhanced adjusted (TFCE; based on 10,000 permutations, http://www.neuro.uni-jena.de/tfce/) and FDR corrected statistics to account for cluster size and multiple comparisons. These whole-brain contrasts were used to first identify clusters of interest in aVS and vmPFC for further analyses, and second to explore other brain areas. To test whether BOLD in aVS and vmPFC was consistent with SV, we first performed a whole-brain conjunction analysis to identify voxels where BOLD signal was higher both for conscious reward (high > low) and subliminal probability (high > low). The aVS cluster of interest was the result of doing a whole-brain analysis with an additional participant excluded because of the lack of signal in ventral aVS (resulting in a larger aVS cluster of interest). We then used the mean beta difference across voxels within the conjunction clusters in aVS and vmPFC to make comparisons between conscious reward, subliminal probability, and subliminal salience.

##### Multivariate pattern similarity analysis

To test for pattern similarities, we used Fischer’s transformed Pearson’s correlations between (e.g., reward and probability) contrast (beta) differences across voxels in our clusters of interest in aVS and vmPFC, and conjunction clusters from the exploratory whole-brain results. Moreover, we used CoSMoMVPA Toolbox (Oosterhof et al., 2016) to perform searchlight procedures with spheres of approx. 30 voxels to test for pattern similarity between beta differences of two contrasts for significant but uncorrected p-values for both contrasts (to create more voxels for the searchlight). For example, doing a searchlight analysis to test for pattern similarity between reward (R5 > R2) and probability (P1 > P0) within voxels that had significant but uncorrected group-level statistics for both reward and probability. The resulting r-maps were TFCE adjusted and corrected for multiple comparisons with Monte Carlo simulations with CoSMoMVPA.

##### Brain-behavior relationships

To test for relationships between BOLD signal change and reaction time, we correlated the mean beta difference across aVS and vmPFC with the mean reaction time difference for conscious reward and subliminal probability. We used Spearman’s rank correlations because of the highly skewed reaction time difference for conscious rewards.

## Results

### Behavioral results

#### Attentional blink

T1 (arrow direction) accuracy was moderately high for the behavioral (M = 78%, SE = .008) and fMRI experiment (M = 81%, SE = .007). The proportion of trials where the probability cue was reported unseen (PAS = 1) was moderately high for the behavioral experiment (M_T2-pres._ = 81%, SE _T2-pres._ = .04; M_T2-abs._ = 86%, SE _T2-abs._ = .03) and very high for the fMRI experiment (M_T2-pres._ = 97%, SE _T2-pres._ = .01; M_T2-abs._ = 97%, SE _T2-abs._ = .01). The difference between experiments was due to the pre-fMRI exclusion of participants with > 20% proportion of seen probability cues. Taken together, this shows that T1 difficulty was sufficient to induce a strong attentional blink effect that renders most probability cues unseen.

#### Value-maximizing choice

When the probability cue was present but unseen (PAS = 1), value-maximizing choice (d’) was not greater than chance in the behavioral experiment when using all trials (M_d’_ = 0.06, SE_d’_ = 0.05, t(33) = 1.29, p = .103, one-tailed) or the fMRI experiment (M_d’_ = −0.10, SE_d’_ = 0.03, t(28) = −3.50, p = .99, one-tailed), suggesting that the subliminal probability cues did not have a direct influence on choice. However, we note that the d’ in the fMRI experiment was unexpectadly negative, which may have been a product of conscious strategies.

The criterion (c) revealed a choice bias towards the risky option when the risky reward was high for the behavioral experiment (high reward, M_c_ = −0.30, SE_c_ = 0.11, t(33) = −2.86, p = .007, two-tailed; low reward, M_c_ = 0.03, SE_c_ = 0.10, t(33) = 0.35, p = .729, two-tailed) and fMRI experiment (high reward, M_c_ = −0.83, SE_c_ = 0.09, t(28) = −9.68, p < .001, two-tailed; low reward, M_c_ = −0.09, Se_c_ = 0.10, t(28) = −0.93, p = .361, two-tailed). Moreover, the choice bias towards risky options for high rewards was greater in the fMRI experiment than the behavioral experiment (high reward, t(61) = 3.82, p < .001, two-tailed; low reward, t(61) = 0.91, p = .366, two-tailed).

#### Reaction time

We found that subliminal probability cues influenced reaction time in both experiments. In the behavioral experiment, the 2 (probability 100% or 0%) x 2 (reward 5 or 2 points) x 2 (risky or safe choice) repeated measures ANOVA revealed a subliminal probability x conscious reward cross-over interaction (F(1,33) = 10.23, p = .003) such that reaction time was faster for high than low reward amount when winning was certain and the reverse when losing was certain. There was no choice x reward (F(1,33) = 3.75, p = .062, choice x probability (F(1,33) = 0.71, p = .405, or choice x reward x probability (F(1,33) = 0.13, p = .718 interactions. There was a main effect of choice (F(1,33) = 10.69, p = .003) such that risky choices were faster than safe choices, but no main effect of conscious rewards (F(1,33) = 0.004, p = .952) or subliminal probabilities (F(1,33) = 4.01, p = .053).

In the fMRI experiment there was no subliminal probability x conscious reward interaction (F(1,28) = 0.13, p = .721). Instead, there was a main effect of subliminal probability (F(1,28) = 6.58, p = .016) such that reaction time was slower when winning was certain than when losing was certain. Moreover, there was a choice x reward interaction (F(1,28) = 13.32, p = .001) such that reaction time was faster for high than low rewards when risky choices were made (t(28) = −5.36, p < .001, two-tailed) but not when safe choices were made (t(28) = 1.02, p = .315, two-tailed). There was no choice x probability interaction (F(1,28) = 0.91, p = .348) or choice x reward x probability interaction (F(1,28) = 2.35, p = .136), but there were main effects of reward (F(1,28) = 16.27, p < .001) and choice (F(1,28) = 35.46, p < .001).

Taken together, the reaction time results show an influence of subliminal probability cues in both experiments but expressed differently. The difference between experiments could be due to interactions between subliminal influences and different conscious strategies or tendencies.

### fMRI results

#### aVS and vmPFC results

Since neural SV signals are expected to correlate positively with reward and probability, we looked for voxels where BOLD signal change was greater for both conscious reward (R5 > R2) and subliminal probability (P1 > P0). Our TFCE adjusted and FDR corrected (p < .05, one-tailed) whole-brain results revealed largely distinct reward- and probability-related clusters throughout the brain, but with the hypothesized overlap in the right aVS (k = 31) and vmPFC (k = 18) (among a few other areas; Figure 2-3), consistent with SV integration. Because of our a priori hypotheses we focused on the conjunction clusters in aVS and vmPFC for further analyses (Figure 2) before conducting exploratory analyses (Figure 3).

**Figure 2.**
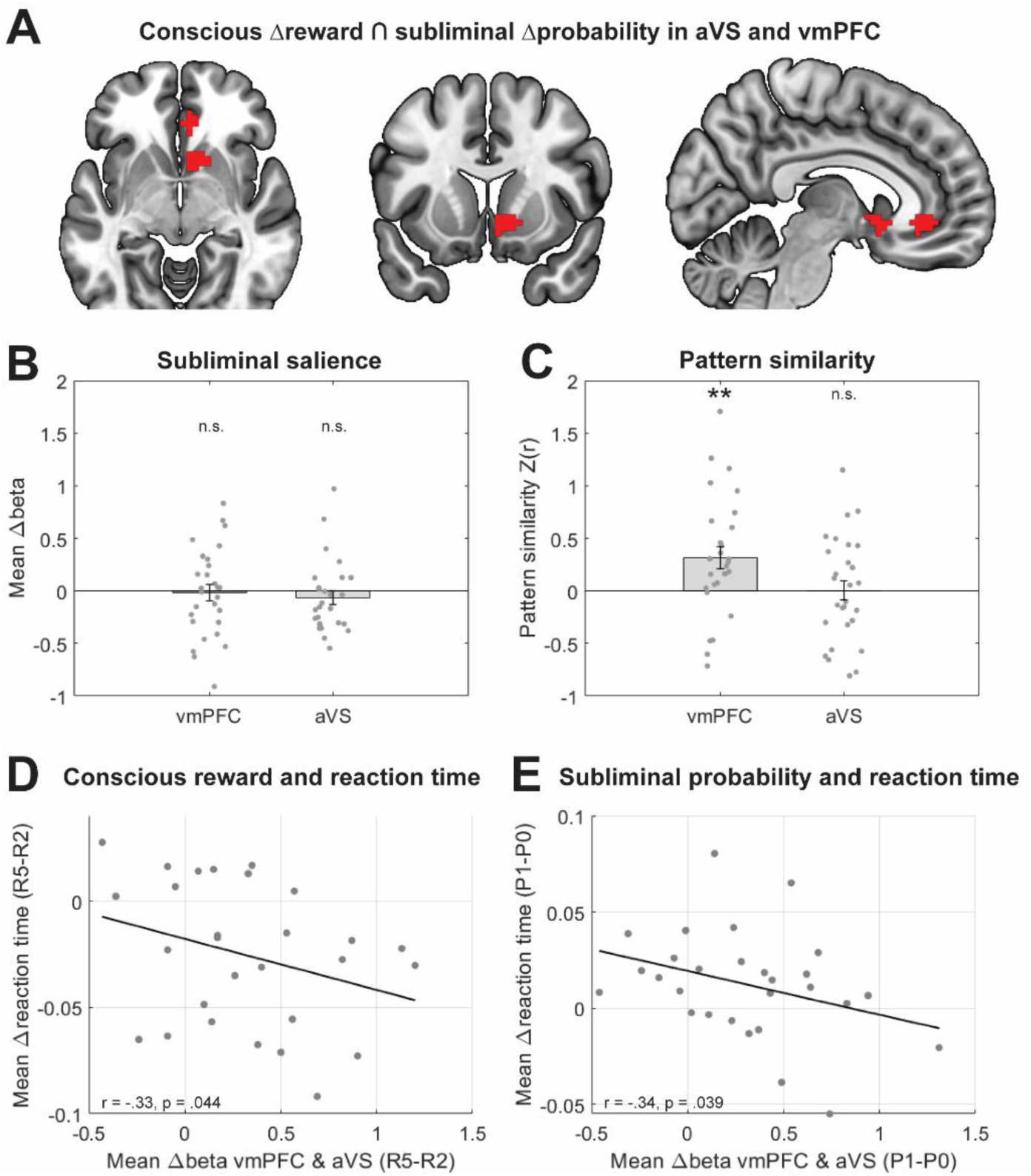
Regions of interest results. (**A**) Conjunction clusters where BOLD signal is greater for conscious reward (R5 > R2) and subliminal probability (P1 > P0) in anterior ventral striatum (aVS) and ventromedial prefrontal cortex (vmPFC). TFCE adjusted and FDR corrected, p < .05, one-tailed at whole-brain level. (**B**) Subliminal salience (cue present > absent) mean beta value difference in aVS and vmPFC clusters. (**C**) Pattern similarity (Fischer’s transformed Pearson’s correlations) between conscious reward and subliminal probability in aVS and vmPFC clusters. (**D**) Spearman’s Rank correlations between mean beta value difference across aVS and vmPFC clusters and mean log10-transformed reaction time difference for conscious reward and subliminal probability, respectively. * p < .05, ** p < .01, n.s. = not significant.

**Figure 3.**
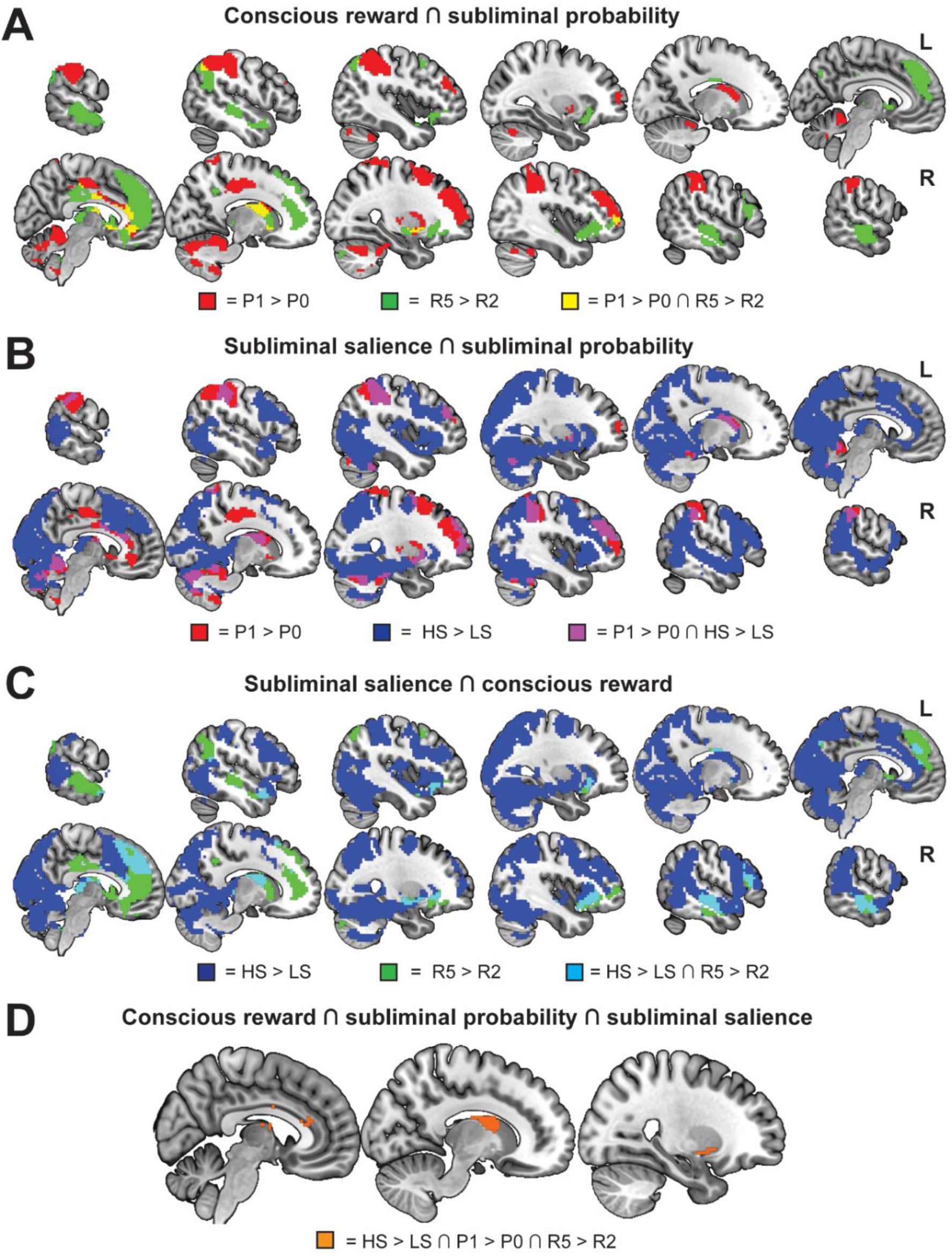
Whole-brain results. Univariate TFCE adjusted and FDR corrected (p < 0.05, one-tailed) contrasts showing conscious reward (green; R5 > R2), subliminal probability (red; P1 > P0), subliminal salience (dark blue; HS > LS), the conjunction of probability and reward (yellow), the conjunction between subliminal salience and subliminal probability (purple), the conjunction of subliminal salience and conscious reward (light blue), and the conjunction between all three contrasts (orange).

Control analyses show that subliminal salience was not greater than zero in aVS (M = −0.65, SE = 0.06, t(27) = −1.01, p = .839, one-tailed) or vmPFC (M = −0.02, SE = 0.08, t(28) = −0.31, p = .620) conjunction clusters (Figure 2B). Moreover, none of the individual voxels in the aVS or vmPFC clusters had significant subliminal salience effects at a liberal p < .05 uncorrected threshold. These results therefore confirm that the conscious reward and subliminal probability effects in aVS and vmPFC cannot be explained by subliminal salience.

If rewards and probabilities are integrated into more abstract “common currency” representations in aVS and vmPFC, then we would expect them to rely on a similar neural code. As expected, we found pattern similarity between conscious rewards and subliminal probabilities within the conjunction cluster in vmPFC (after removing an outlier with negative Z(r) > 3 SD, M_Z(r)_ = 0.32, SE_Z(r)_ = 0.11, t(27) = 2.93, p = .0003, one-tailed; and using all participants, M_Z(r)_ = 0.25, SE_Z(r)_ = 0.12, t(28) = 2.07, p = .024, one-tailed) (Figure 2C). Contrary to our expectations, we did not find pattern similarity between conscious rewards and subliminal probabilities in aVS (M_Z(r)_ = 0.006, SE_Z(r)_ = 0.09, t(27) = 0.06, p = .476, one-tailed). The pattern similarity results suggest that conscious rewards and subliminal probabilities are integrated into a more abstract value representation in vmPFC.

Because previous literature show that choice reaction times are faster for larger SV differences between options (Krajbich et al., 2010), we tested for and found Spearman’s Rank correlations between mean beta differences across aVS and vmPFC clusters and reaction time differences for conscious reward (r(27) = −.328, p = .044, one-tailed; Figure 2C) and subliminal probability (r(27) = −.339 p = .039, one-tailed; Figure 2D), linking the neural signal in putative SV areas with reaction time in a manner consistent with SV.

#### Whole-brain results

The three whole-brain contrasts for (high > low) conscious rewards, subliminal probabilities, and subliminal salience revealed largely distinct brain activations with some overlap between them (Figure 3; TFCE adjusted and FDR corrected, p < .05, one-tailed). Conscious rewards (R5 > R2) were associated with BOLD signal change in bilateral dmPFC extending to the right vmPFC; bilateral aVS, aINS/frontal orbital cortex, thalamus, brainstem, and superior/middle temporal cortex; right caudate nucleus, putamen, and inferior frontal cortex (S-Table 1). Subliminal probabilities (P1 > P0) were associated with BOLD signal change in right vmPFC, aVS, PCC, and middle/superior frontal gyrus; bilateral caudate nucleus, putamen, frontal poles, supra marginal gyrus (SMG), superior parietal lobe (SPL), and cerebellum (S-Table 2). Subliminal salience (unseen probability cue present > absent) was associated with BOLD signal change in bilateral dmPFC extending to left vmPFC/subcallosal cortex; bilateral occipital and ventral temporal cortex, middle/superior temporal cortex and temporal poles, posterior parietal cortex (SMG/AG/SPL), aINS/orbitofrontal cortex, caudate nucleus, putamen, hippocampus, amygdala, thalamus, brainstem, and precentral/middle frontal gyrus (S-Table 3).

In addition to aVS and vmPFC, there was overlapping BOLD signal change between conscious rewards and subliminal probabilities in the right caudate nucleus, right dorsal anterior cingulate cortex (dACC), right frontal pole, left AG, right putamen, right superior frontal gyrus, right precentral gyrus, mid/posterior cingulate cortex (Figure 3A; S-Table 4). Moreover, there was overlapping BOLD signal change between subliminal probability and salience in bilateral cerebellum, SMG, prefrontal/frontal pole, putamen, caudate nucleus, and dACC; right middle/superior frontal gyrus, amygdala, and superior parietal lobe (Figure 3B; S-Table 5). There was overlapping BOLD signal change between conscious reward and subliminal salience in bilateral aINS/frontal orbital cortex, dmPFC, middle temporal gyrus, temporal pole, cerebellum, brainstem, hippocampus, and amygdala; left vmPFC/subcallosal cortex and SMG/AG; and right caudate nucleus (Figure 3C; S-Table 6). Lastly, there were three brain areas in which BOLD signal change overlapped for all three contrasts: right caudate nucleus, putamen, and dACC (Figure 3D; S-Table 7).

When testing pattern similarity between reward and probability, salience and probability, and salience and reward in the respective overlap areas with a cluster size of at least ten voxels none of the areas survived FDR corrections. Additionally, we performed a pattern similarity searchlight analysis between the three contrasts, but nothing survived corrections for multiple comparison (though the most prominent cluster of all uncorrected searchlight similarity results was between conscious reward and subliminal probability in the right vmPFC overlapping our cluster of interest in Figure 2).

## Discussion

Here we investigated whether subliminal probabilities were integrated with conscious rewards to form abstract SV representations in aVS and vmPFC. We found (i) that BOLD signal change was greater for conscious rewards (high > low) and subliminal probabilities (high > low) in aVS and vmPFC, which is consistent with neural SV signals; (ii) that neither the conscious reward nor the subliminal probability effects in aVS or vmPFC could be explained by salience, (iii) that subliminal probabilities and conscious rewards had similar neural patterns in vmPFC but not in aVS; and (iv) that mean neural signals related to conscious rewards and subliminal probabilities in aVS and vmPFC correlated negatively with reaction time. Taken together, these findings suggest that conscious awareness is not necessary for subliminal probabilities to be integrated with conscious rewards to form a more abstract “common currency” SV representation in vmPFC. Additionally, exploratory whole-brain contrasts revealed brain-wide BOLD signal change related to subliminal probabilities and salience, suggesting that information can have “global access” without conscious awareness.

### Subjective value integration without conscious awareness

As expected, we found that BOLD signal change was greater for high than low conscious rewards and high than low subliminal probabilities in aVS and vmPFC, which could not be explained by subliminal salience. Moreover, the multivariate pattern similarity analyses revealed similar neural patterns for conscious rewards and subliminal probabilities in vmPFC, suggesting that they were integrated into more abstract shared value representations. Our findings are thus consistent with literature showing that SV (but not salience) correlates positively with BOLD signal in aVS and vmPFC across tasks, stimuli, and domains, and is therefore thought to represent a more abstract domain-general “common currency” representation (Bartra et al., 2013; Chib et al., 2009; Dang et al., 2024; Kable & Glimcher, 2007, 2009; Kobayashi & Hsu, 2019; D. J. Levy & Glimcher, 2012; I. Levy et al., 2010; McNamee et al., 2013; Pegors et al., 2015; Shuster & Levy, 2018). Consistent with these findings, individual differences in anatomical structure or lesions to vmPFC are associated with differences or disruptions in choice preferences (Bergström, Lerman, et al., 2024; Bergström, Schu, et al., 2024; Camille et al., 2011; Fellows & Farah, 2007; Yu et al., 2022). The idea of a domain-general “common currency” has mainly been supported by overlapping univariate results with a few studies using multivariate techniques to demonstrate pattern similarity of SV across domains, such as visual and auditory modalities (Dang et al., 2024; Shuster & Levy, 2018), information and basic rewards (Kobayashi & Hsu, 2019), food, money, and trinkets (McNamee et al., 2013), and face and place attractiveness (Pegors et al., 2015). Adding to previous findings, our novel results demonstrate pattern similarity across awareness (conscious and subliminal) and value modality (reward and probability). Future work will have to address whether pattern similarity across conscious probabilities and rewards consistently show an effect in vmPFC but not aVS, which would suggest a difference in abstraction between the two areas. Consistent with literature showing faster reaction time for higher SV (Krajbich et al., 2010), we also found that mean BOLD signal change across putative SV areas (aVS and vmPFC) correlated negatively with reaction time, despite the main effect of reaction time being slower for high than low subliminal probability. A likely explanation is that the final reaction time may have many different conscious and/or subliminal influences other than SV. Taken together, these results suggest that subliminal probabilities were integrated with conscious rewards in aVS and vmPFC to form a more abstract neural SV representation in vmPFC.

Importantly, our results challenge the idea that subliminal value-related information is confined to subcortical processing (Bijleveld et al., 2012b). Our results also add to previous findings that subliminal rewards are processed subcortically (Pessiglione et al., 2007, 2008), by showing that subliminal probabilities also are processed in similar subcortical areas. The subliminal instrumental learning result found in Pessiglione et al. (2008) has been challenged on account of not being replicated with trial-wise subjective measures of awareness, suggesting that those results may have been influenced by conscious awareness (Skora et al., 2021, 2023), which then also calls into question the associations with BOLD signal in aVS. However, the fact that Pessiglione et al. (2008) was not replicated with trial-wise measures of awareness does not necessarily mean that their results were driven by conscious influences. Changing to trial-wise awareness measures would likely lower the threshold for conscious awareness by shifting attentional resources to the otherwise subliminal stimuli, potentially leading to shorter stimuli durations in staircase procedures or discarded trials based on awareness measures, and therefore weaker subliminal effects. Nevertheless, here we used trial-wise measures of conscious awareness to be able to discard individual trials with seen probability cues to avoid such criticism. Moreover, the information integration in vmPFC adds to the wider literature on integration of subliminal information (for review, see Mudrik et al., 2014). Although we show subliminal integration of a subliminal and a conscious component, future research will have to explore whether two subliminal value-components (e.g., reward and probability) also can be integrated.

### Subliminal salience

Though no subliminal salience effects were found in aVS or vmPFC, there were brain-wide salience effects in many other areas previously associated with (conscious) salience, such as dmPFC, PPC, aINS, and caudate nucleus (Bartra et al., 2013; Kahnt & Tobler, 2017; Litt et al., 2011; Zink et al., 2004). However, we did not find any evidence of abstract salience representations when testing for pattern similarity between salience and reward or probability. This was also true in the right caudate nucleus and putamen areas with overlapping univariate results for salience, probability, and reward, suggesting these areas integrate this information without relying on a shared neural code to do so. Our salience effects are from learned associations of certainty (100% and 0% to win) compared to uncertainty (50% to win), while holding average BOLD signal change related to rewards and probabilities constant. Taken together, we thus demonstrate the presence of neural effects from subliminal top-down salience.

### Global access without conscious awareness

Interestingly, the brain-wide effects of both subliminal salience and subliminal probability suggests that information can have “global access” without conscious awareness. The GNW theory posits that subliminal information is processed in parallel in localized modular circuitry that becomes conscious only if given global access via a “workspace” of thalamocortical and frontoparietal network (Dehaene & Changeux, 2011; Dehaene & Naccache, 2001; Mashour et al., 2020; Melloni et al., 2023). Information is thought to remain subliminal if too weak to enter the global workspace or if the global workspace is occupied by other conscious information (Melloni et al., 2023). In attentional blink paradigms, one could argue that information likely is strong enough to enter but that the workspace is occupied by other information (i.e., the T1 task and distractors). Our findings suggest that subliminal information can have global access in the sense that it can spread through many different localized modules or parts of the brain at multiple hierarchical levels, including frontoparietal areas. However, it remains unclear whether the subliminal information spread via the frontoparietal network or through other means. Future research will have to explore how subliminal information can have this kind of global access.

### Subliminal behavioral influences

In both experiments, subliminal probability cues influenced reaction time despite value-maximizing choice (d’) not being greater than chance. Previous literature on reaction times when choice information was conscious would have predicted main effects of conscious rewards and subliminal probabilities such that reaction time was faster for higher than lower reward and probability values (Krajbich et al., 2010). However, that was not the case in either of the two experiments. In the behavioral experiment we found a subliminal probability x conscious reward interaction, such that reaction time was faster for high than low reward when winning was certain and the reverse when losing was certain. By contrast, the fMRI experiment revealed a choice x reward interaction such that reaction time was faster for high than low reward only when risky choices were made, and a main effect of subliminal probabilities such that reaction time was slower for high than low probability. It is likely that the subliminal nature of the probability cues substantially changed the task and how participants responded to it, and introduced different conscious strategies that may interact with the subliminal information, to cause different influences on reaction time and value-maximizing choice.

### Subliminal memory retention

The fact that the subliminal probability cues influenced reaction time without being erased by subsequent distractors suggests some kind of robust subliminal memory retention. We cannot make any claims or add to the debate about subliminal instrumental learning (Pessiglione et al., 2008; Skora et al., 2021, 2023) because our participants consciously learned the associations between cue and probability prior to the experiments. However, contrary to claims that conscious awareness is necessary for any kind of instrumental reactivation to previously learned information (Skora et al., 2024), our findings do show that consciously learned associations can be subliminally reactivated with neural and reaction time influences. Additionally, we can infer that cue retention in our experiments are not related to sensory/iconic memory mechanisms as the distractors would have overwritten such residual memory traces (Pinto et al., 2013), and not to repetition priming as there was no information repetition. The salience contrast (cue present > absent) revealed BOLD signal change in cerebellum, hippocampus, visual cortex, and frontoparietal areas, suggesting that subliminal cue information could have been reactivated and/or retained via hippocampus-(Degonda et al., 2005; Henke et al., 2003; Reber et al., 2012), cerebellum-based associations, and/or working memory maintenance (Bergström & Eriksson, 2014, 2015, 2018; King et al., 2016; Pedale et al., 2023; Soto et al., 2011).

## Conclusions

In sum, we show that conscious awareness is not necessary for probabilities to be integrated with conscious rewards to form SV representations in the vmPFC. Moreover, the brain-wide subliminal probability and salience effects suggest that information can have global access without conscious awareness. Future research will have to further explore the relevance and impact of conscious awareness on different value computations during choice.

## Author Contributions

FB: Conceptualization, funding acquisition, supervision, project administration, methodology, investigation, formal analysis, visualization, validation, data curation, writing – original draft, writing – review & editing. PF: Investigation, formal analysis, writing – original draft, writing – review & editing. SL: Methodology and writing – review & editing JWK: Methodology and writing – review & editing. PS: Methodology and writing – review & editing. JA: Methodology and writing – review & editing. JE: Methodology and writing – review & editing. BS: Methodology and writing – review & editing

## Declaration of Competing Interest

The authors declare no competing financial interests.

## Funding/Acknowledgements

This work was supported by The Bial Foundation (A-27942 to FB). FB was supported by Fundação para a Ciência e Tecnologia (CEECIND/03661/2017). JA was supported by a European Research Counsil (ERC) under the European Union’s Horizon 2020 research and innovation program Starting Grant number 802553 “Content Map”, and by European Research Executive Agency Widening program under the European Union’s Horizon Europe Grant 101087584 “Cogbooster”. We thank Alice Pacheco for assisting with data collection for the behavioral experiment.

## Notes

### Competing Interest Statement

The authors have declared no competing interest.

