## Supplemental Material for "Subliminal risk influences subjective value in the ventromedial prefrontal cortex"

#### Method

##### fMRI preprocessing

Results included in this manuscript come from preprocessing performed using *fMRIPrep* 23.2.0a2 (Esteban et al. (2019); Esteban et al. (2018); RRID:SCR_016216), which is based on *Nipype* 1.8.6 (K. Gorgolewski et al. (2011); K. J. Gorgolewski et al. (2018); RRID:SCR_002502).

*Anatomical data preprocessing*. A total of 1 T1-weighted (T1w) images were found within the input BIDS dataset. The T1w image was corrected for intensity non-uniformity (INU) with *N4BiasFieldCorrection* (Tustison et al. 2010), distributed with ANTs 2.5.0 (Avants et al. 2008, RRID:SCR_004757), and used as T1w-reference throughout the workflow. The T1w-reference was then skull-stripped with a *Nipype* implementation of the *antsBrainExtraction.sh* workflow (from ANTs), using OASIS30ANTs as target template. Brain tissue segmentation of cerebrospinal fluid (CSF), white-matter (WM) and gray-matter (GM) was performed on the brain-extracted T1w using fast (FSL (version unknown), RRID:SCR_002823, Zhang, Brady, and Smith 2001). Brain surfaces were reconstructed using recon-all (FreeSurfer 7.3.2, RRID:SCR_001847, Dale, Fischl, and Sereno 1999), and the brain mask estimated previously was refined with a custom variation of the method to reconcile ANTs-derived and FreeSurfer-derived segmentations of the cortical gray-matter of *Mindboggle* (RRID:SCR_002438, Klein et al. 2017). Volume-based spatial normalization to one standard space (MNI152NLin2009cAsym) was performed through nonlinear registration with *antsRegistration* (ANTs 2.5.0), using brain-extracted versions of both T1w reference and the T1w template. The following template was selected for spatial normalization and accessed with *TemplateFlow* (23.1.0, Ciric et al. 2022): *ICBM 152 Nonlinear Asymmetrical template version 2009c* [Fonov et al. (2009), RRID:SCR_008796; TemplateFlow ID: MNI152NLin2009cAsym].

*Functional data preprocessing*. For each of the 4 BOLD runs found per subject (across all tasks and sessions), the following preprocessing was performed. First, a reference volume was generated, using a custom methodology of *fMRIPrep*, for use in head motion correction. Head-motion parameters with respect to the BOLD reference (transformation matrices, and six corresponding rotation and translation parameters) are estimated before any spatiotemporal filtering using *mcflirt* (FSL). The BOLD reference was then co-registered to the T1w reference using *bbregister* (FreeSurfer) which implements boundary-based registration (Greve and Fischl 2009). Co-registration was configured with six degrees of freedom. Several confounding time-series were calculated based on the preprocessed BOLD: framewise displacement (FD), DVARS and three region-wise global signals. FD was computed using two formulations following Power (absolute sum of relative motions, Power et al. (2014)) and Jenkinson (relative root mean square displacement between affines, Jenkinson et al. (2002)). FD and DVARS are calculated for each functional run, both using their implementations in *Nipype* (following the definitions by Power et al. 2014). The three global signals are extracted within the CSF, the WM, and the whole-brain masks. Additionally, a set of physiological regressors were extracted to allow for component-based noise correction (*CompCor*; Behzadi, Restom, Liau, & Liu, 2007). Principal components are estimated after high-pass filtering the preprocessed BOLD time-series (using a discrete cosine filter with 128s cut-off) for the two *CompCor* variants: temporal (*tCompCor*) and anatomical (*aCompCor*). *tCompCor* components are then calculated from the top 2% variable voxels within the brain mask. For *aCompCor*, three probabilistic masks (CSF, WM and combined CSF + WM) are generated in anatomical space. The implementation differs from that of Behzadi et al. in that instead of eroding the masks by 2 pixels on BOLD space, a mask of pixels that likely contain a volume fraction of GM is subtracted from the *aCompCor* masks. This mask is obtained by dilating a GM mask extracted from the FreeSurfer's *aseg* segmentation, and it ensures components are not extracted from voxels containing a minimal fraction of GM. Finally, these masks are resampled into BOLD space and binarized by thresholding at 0.99 (as in the original implementation). Components are also calculated separately within the WM and CSF masks. For each CompCor decomposition, the k components with the largest singular values are retained, such that the retained components' time series are sufficient to explain 50 percent of variance across the nuisance mask (CSF, WM, combined, or temporal). The remaining components are dropped from consideration. The head-motion estimates calculated in the correction step were also placed within the corresponding confounds file. The confound time series derived from head motion estimates and global signals were expanded with the inclusion of temporal derivatives and quadratic terms for each (Satterthwaite et al. 2013). Frames that exceeded a threshold of 0.5 mm FD or 1.5 standardized DVARS were annotated as motion outliers. Additional nuisance time-series are calculated by means of principal components analysis of the signal found within a thin band (crown) of voxels around the edge of the brain, as proposed by (Patriat, Reynolds, and Birn 2017). All resamplings can be performed with a single interpolation step by composing all the pertinent transformations (i.e. head-motion transform matrices, susceptibility distortion correction when available, and co-registrations to anatomical and output spaces). Gridded (volumetric) resamplings were performed using *nitransforms*, configured with cubic B-spline interpolation.

Many internal operations of *fMRIPrep* use *Nilearn* 0.10.2 (Abraham et al. 2014, RRID:SCR_001362), mostly within the functional processing workflow. For more details of the pipeline, see the section corresponding to workflows in *fMRIPrep's* documentation.

*Copyright Waiver*. The above boilerplate text was automatically generated by fMRIPrep with the express intention that users should copy and paste this text into their manuscripts unchanged. It is released under the CC0 license.

### Results

**S-Table 1.**
*Conscious reward (R5 > R2)*

| Brain area | Hemi. | MNI | TFCE | k |
|---|---|---|---|---|
| Dorso- & ventromedial prefrontal cortex, and anterior ventral striatum | B | [5 45 33] | 1790 | 1306 |
| Middle & superior temporal gyrus | L | [-55 -27 -6] | 538 | 432 |
| Anterior insula & orbitofrontal cortex | R | [35 24 -15] | 558 | 399 |
| Middle & superior temporal gyrus, putamen, and amygdala | R | [46 -15 -9] | 251 | 273 |
| Inferior parietal lobule | L | [-55 -57 36] | 271 | 163 |
| Posterior cingulate cortex | B | [2 -27 36] | 296 | 122 |

| | | | | |
|---|---|---|---|---|
| Cerebellum crus 1 | R | [20 -75 -33] | 97.5 | 19 |
| Precuneus & posterior cingulate | R | [13 -48 39] | 119 | 16 |
| Middle frontal gyrus | L | [-44 12 51] | 91.9 | 15 |
| Frontal pole | L | [-21 57 18] | 112 | 12 |
| Amygdala & hippocampus | R | [16 -12 -12] | 159 | 10 |
| Frontal pole | R | [16 60 12] | 83.8 | 10 |
| Precuneus | L | [-6 -69 36] | 101 | 10 |
| Pre/post central | R | [20 -30 63] | 103 | 8 |
| Ventromedial prefrontal cortex | R | [5 42 -24] | 90.6 | 6 |

Note. The table shows results for threshold-free cluster enhanced (TFCE) FDR corrected results at p < .05 with names of brain areas, peak voxel MNI coordinate, peak voxel statistical TFCE value and cluster size (k). Clusters with k < 5 are omitted for brevity.

**S-Table 2.**

*Subliminal probability (P1 > P0)*

| Brain area | Hemi. | MNI | TFCE | k |
|---|---|---|---|---|
| Superior & middle frontal gyrus, and frontal pole | R | [28 9 63] | 293 | 1182 |
| Cerebellum I-V | B | [5 -45 -18] | 442 | 627 |
| Supramarginal gyrus & angular gyrus | L | [-47 -48 51] | 626 | 611 |
| Ventromedial PFC, anterior & posterior cingulate | R | [13 -30 39] | 235 | 227 |
| Caudate nucleus & putamen | R | [16 -3 18] | 235 | 220 |
| Caudate nucleus & putamen | L | [-17 6 12] | 180 | 102 |
| Frontal pole | L | [-36 54 18] | 156 | 60 |
| Cerebellum Crus I | L | [-44 -48 -36] | 179 | 27 |
| Cerebellum I-V | L | [-17 -42 -21] | 167 | 23 |
| Anterior ventral striatum | R | [9 9 -6] | 163 | 23 |
| Cerebellum Crus I & VI | L | [-28 -66 -30] | 173 | 22 |
| Cerebellum Crus I | L | [-44 -75 -30] | 165 | 16 |
| Orbitofrontal cortex | L | [-25 24 -15] | 121 | 6 |

Note. The table shows results for threshold-free cluster enhanced (TFCE) FDR corrected results at p < .05 with names of brain areas, peak voxel MNI coordinate, peak voxel statistical TFCE value and cluster size (k). Clusters with k < 5 are omitted for brevity.

**S-Table 3.**

*Subliminal Salience (cue present > absent)*

| Brain area | Hemi. | MNI | TFCE | k |
|---|---|---|---|---|
| Occipital cortex | B | [-17 -69 9] | 2450 | 17180 |
| Ventral temporal cortex | B | | | |
| Middle temporal cortex | B | | | |
| Superior temporal cortex | B | | | |
| Temporal poles | B | | | |
| Inferior / superior parietal lobule | B | | | |

| | |
|---|---|
| Anterior insula / orbitofrontal cortex | B |
| Caudate nucleus | B |
| Putamen | B |
| Hippocampus | B |
| Amygdala | B |
| Brainstem | B |
| Precentral / middle frontal gyrus | B |

Note. The table shows results for threshold-free cluster enhanced (TFCE) FDR corrected results at p < .05 with names of brain areas, peak voxel MNI coordinate, peak voxel statistical TFCE value and cluster size (k). Clusters with k < 5 are omitted for brevity.

**S-Table 4.**

*Conscious reward (R5 > R2) ∩ Subliminal probability (P1 > P0)*

| **Brain area** | **Hemi.** | **MNI** | **TFCE** | **k** |
|---|---|---|---|---|
| Caudate nucleus and Thalamus | R | [13 -3 15] | 333 | 61 |
| Anterior ventral striatum | R | [9 9 -3] | 357 | 21 |
| Anterior cingulate cortex | R | [9 36 15] | 404 | 20 |
| Frontal pole | R | [35 51 9] | 159 | 19 |
| ventromedial prefrontal cortex | R | [5 39 -3] | 250 | 18 |
| Inferior parietal lobule | L | [-55 -60 45] | 185 | 18 |
| Putamen | R | [32 -6 -6] | 183 | 14 |
| Superior frontal gyrus | R | [20 30 48] | 203 | 12 |
| Pre and post central gyrus | R | [20 -30 63] | 144 | 8 |
| Middle cingulate gyrus | R | [5 -9 33] | 138 | 6 |

Note. The table shows results for threshold-free cluster enhanced (TFCE) FDR corrected results at p < .05 with names of brain areas, peak voxel MNI coordinate, peak voxel statistical TFCE value (mean across conjunctive results) and cluster size (k). Clusters with k < 5 are omitted for brevity.

**S-Table 5.**

*Subliminal salience (cue present > absent) ∩ Subliminal probability (P1 > P0)*

| **Brain area** | **Hemi.** | **MNI** | **TFCE** | **k** |
|---|---|---|---|---|
| Cerebellum, Crus I, I-IV, V | R | [13 -72 -24] | 730 | 391 |
| Inferior and superior parietal lobule | L | [-44 -42 48] | 701 | 282 |
| Frontal pole and middle frontal gyrus | R | [32 51 30] | 344 | 253 |
| Inferior and superior parietal lobule | R | [39 -45 42] | 434 | 225 |
| Caudate nucleus and Putamen | L | [-25 15 6] | 308 | 92 |
| Superior and middle frontal gyrus | R | [35 3 45] | 344 | 56 |
| Caudate and Thalamus | R | [16 -3 18] | 294 | 54 |
| Putamen | R | [24 9 -3] | 240 | 39 |
| Cerebellum Crus I and Crus II | L | [-40 -48 -33] | 392 | 27 |
| Superior parietal lobule | R | [9 -54 72] | 352 | 27 |
| Anterior cingulate cortex | R | [-2 24 21] | 251 | 25 |
| Cerebellum VI and Crus I | L | [-25 -69 -27] | 692 | 22 |

| | | | | |
|---|---|---|---|---|
| Frontal pole and middle frontal gyrus | L | [-47 39 24] | 239 | 22 |
| Putamen | R | [32 -12 -3] | 209 | 8 |
| Cerebellum Crus I | L | [-40 -78 -24] | 236 | 6 |
| Anterior cingulate cortex | R | [5 3 30] | 239 | 6 |
| Cerebellum I-IV and V | L | [-17 -36 -21] | 150 | 5 |

Note. The table shows results for threshold-free cluster enhanced (TFCE) FDR corrected results at p < .05 with names of brain areas, peak voxel MNI coordinate, peak voxel statistical TFCE value (mean across conjunctive results) and cluster size (k). Clusters with k < 5 are omitted for brevity.

**S-Table 6.**

*Subliminal salience (cue present > absent) ∩ conscious reward (R5 >R2)*

| Brain area | Hemi. | MNI | TFCE | k |
|---|---|---|---|---|
| Anterior insula and orbitofrontal cortex, and inferior and middle frontal gyrus | R | [43 18 27] | 512 | 289 |
| Dorsomedial prefrontal cortex | R | [5 45 33] | 1040 | 283 |
| Superior and middle temporal gyrus, Putamen and amygdala | R | [58 -27 -6] | 484 | 217 |
| Thalamus and Caudate | R | [9 -30 3] | 744 | 142 |
| Temporal pole | L | [-55 6 -21] | 342 | 140 |
| Temporoparietal junction | L | [-55 -54 18] | 182 | 15 |
| Subcallosal cortex | B | [2 27 -12] | 172 | 11 |
| Cerebellum Crus I | R | [20 -78 -30] | 249 | 9 |
| Hippocampus and amygdala | R | [16 -9 -12] | 373 | 9 |
| Brainstem | B | [2 -21 -18] | 73.5 | 7 |

Note. The table shows results for threshold-free cluster enhanced (TFCE) FDR corrected results at p < .05 with names of brain areas, peak voxel MNI coordinate, peak voxel statistical TFCE value (mean across conjunctive results) and cluster size (k). Clusters with k < 5 are omitted for brevity.

**S-Table 7.**

*Subliminal salience (cue present > absent) ∩ subliminal probability (P1 > P0) ∩ conscious reward (R5 > R2)*

| Brain area | Hemi. | MNI | TFCE | k |
|---|---|---|---|---|
| Caudate nucleus | R | [13 -3 15] | 339 | 50 |
| Putamen | R | [24 6 -6] | 187 | 10 |
| Anterior cingulate cortex | R | [5 33 18] | 267 | 7 |

Note. The table shows results for threshold-free cluster enhanced (TFCE) FDR corrected results at p < .05 with names of brain areas, peak voxel MNI coordinate, peak voxel statistical TFCE value (mean across conjunctive results) and cluster size (k). Clusters with k < 5 are omitted for brevity.

#### References

Abraham, Alexandre, Fabian Pedregosa, Michael Eickenberg, Philippe Gervais, Andreas Mueller, Jean Kossaifi, Alexandre Gramfort, Bertrand Thirion, and Gael Varoquaux. 2014. “Machine

Learning for Neuroimaging with Scikit-Learn." Frontiers in Neuroinformatics 8. https://doi.org/10.3389/fninf.2014.00014.

Avants, B. B., C. L. Epstein, M. Grossman, and J. C. Gee. 2008. "Symmetric Diffeomorphic Image Registration with Cross-Correlation: Evaluating Automated Labeling of Elderly and Neurodegenerative Brain." Medical Image Analysis 12 (1): 26–41. https://doi.org/10.1016/j.media.2007.06.004.

Behzadi, Yashar, Khaled Restom, Joy Liau, and Thomas T. Liu. 2007. "A Component Based Noise Correction Method (CompCor) for BOLD and Perfusion Based fMRI." NeuroImage 37 (1): 90–101. https://doi.org/10.1016/j.neuroimage.2007.04.042.

Ciric, R., William H. Thompson, R. Lorenz, M. Goncalves, E. MacNicol, C. J. Markiewicz, Y. O. Halchenko, et al. 2022. "TemplateFlow: FAIR-Sharing of Multi-Scale, Multi-Species Brain Models." Nature Methods 19: 1568–71. https://doi.org/10.1038/s41592-022-01681-2.

Dale, Anders M., Bruce Fischl, and Martin I. Sereno. 1999. "Cortical Surface-Based Analysis: I. Segmentation and Surface Reconstruction." NeuroImage 9 (2): 179–94. https://doi.org/10.1006/nimg.1998.0395.

Esteban, Oscar, Ross Blair, Christopher J. Markiewicz, Shoshana L. Berleant, Craig Moodie, Feilong Ma, Ayse Ilkay Isik, et al. 2018. "fMRIPrep 23.2.0a2." Software. https://doi.org/10.5281/zenodo.852659.

Esteban, Oscar, Christopher Markiewicz, Ross W Blair, Craig Moodie, Ayse Ilkay Isik, Asier Erramuzpe Aliaga, James Kent, et al. 2019. "fMRIPrep: A Robust Preprocessing Pipeline for Functional MRI." Nature Methods 16: 111–16. https://doi.org/10.1038/s41592-018-0235-4.

Fonov, VS, AC Evans, RC McKinstry, CR Almli, and DL Collins. 2009. "Unbiased Nonlinear Average Age-Appropriate Brain Templates from Birth to Adulthood." NeuroImage 47, Supplement 1: S102. https://doi.org/10.1016/S1053-8119(09)70884-5.

Gorgolewski, K., C. D. Burns, C. Madison, D. Clark, Y. O. Halchenko, M. L. Waskom, and S. Ghosh. 2011. "Nipype: A Flexible, Lightweight and Extensible Neuroimaging Data Processing Framework in Python." Frontiers in Neuroinformatics 5: 13. https://doi.org/10.3389/fninf.2011.00013.

Gorgolewski, Krzysztof J., Oscar Esteban, Christopher J. Markiewicz, Erik Ziegler, David Gage Ellis, Michael Philipp Notter, Dorota Jarecka, et al. 2018. "Nipype." Software. https://doi.org/10.5281/zenodo.596855.

Greve, Douglas N, and Bruce Fischl. 2009. "Accurate and Robust Brain Image Alignment Using Boundary-Based Registration." NeuroImage 48 (1): 63–72. https://doi.org/10.1016/j.neuroimage.2009.06.060.

Jenkinson, Mark, Peter Bannister, Michael Brady, and Stephen Smith. 2002. "Improved Optimization for the Robust and Accurate Linear Registration and Motion Correction of Brain Images." NeuroImage 17 (2): 825–41. https://doi.org/10.1006/nimg.2002.1132.

Klein, Arno, Satrajit S. Ghosh, Forrest S. Bao, Joachim Giard, Yrjö Häme, Eliezer Stavsky, Noah Lee, et al. 2017. "Mindboggling Morphometry of Human Brains." PLOS Computational Biology 13 (2): e1005350. https://doi.org/10.1371/journal.pcbi.1005350.

Patriat, Rémi, Richard C. Reynolds, and Rasmus M. Birn. 2017. "An Improved Model of Motion-Related Signal Changes in fMRI." NeuroImage 144, Part A (January): 74–82. https://doi.org/10.1016/j.neuroimage.2016.08.051.

Power, Jonathan D., Anish Mitra, Timothy O. Laumann, Abraham Z. Snyder, Bradley L. Schlaggar, and Steven E. Petersen. 2014. “Methods to Detect, Characterize, and Remove Motion Artifact in Resting State fMRI.” NeuroImage 84 (Supplement C): 320–41. https://doi.org/10.1016/j.neuroimage.2013.08.048.

Satterthwaite, Theodore D., Mark A. Elliott, Raphael T. Gerraty, Kosha Ruparel, James Loughead, Monica E. Calkins, Simon B. Eickhoff, et al. 2013. “An improved framework for confound regression and filtering for control of motion artifact in the preprocessing of resting-state functional connectivity data.” NeuroImage 64 (1): 240–56. https://doi.org/10.1016/j.neuroimage.2012.08.052.

Tustison, N. J., B. B. Avants, P. A. Cook, Y. Zheng, A. Egan, P. A. Yushkevich, and J. C. Gee. 2010. “N4ITK: Improved N3 Bias Correction.” IEEE Transactions on Medical Imaging 29 (6): 1310–20. https://doi.org/10.1109/TMI.2010.2046908.

Zhang, Y., M. Brady, and S. Smith. 2001. “Segmentation of Brain MR Images Through a Hidden Markov Random Field Model and the Expectation-Maximization Algorithm.” IEEE Transactions on Medical Imaging 20 (1): 45–57. https://doi.org/10.1109/42.906424.
